# Heme stimulates platelet mitochondrial oxidant production via the activation of toll-like receptor 4 signaling to mediate targeted granule secretion

**DOI:** 10.1101/2021.08.02.454816

**Authors:** Gowtham K. Annarapu, Deirdre Nolfi-Donegan, Michael Reynolds, Yinna Wang, Lauren Kohut, Brian Zuckerbraun, Sruti Shiva

## Abstract

Hemolysis is a pathological component of many diseases and is associated with thrombosis and vascular dysfunction. Hemolytic products, including cell-free hemoglobin and free heme directly activate platelets. However, the effect of hemolysis on platelet degranulation, a central process in not only thrombosis, but also inflammatory and mitogenic signaling, remains less clear. Our group showed that hemoglobin-induced platelet activation involved the production of mitochondrial reactive oxygen species (mtROS). However, the molecular mechanism by which extracellular hemolysis induces platelet mtROS production, and whether the mtROS regulate platelet degranulation remains unknown. Here, we demonstrate using isolated human platelets that cell free heme is a more potent agonist for platelet activation than hemoglobin, and stimulates the release of a specific set of molecules from the α-granule of platelets, including the glycoprotein thrombospondin-1 (TSP-1). We uncover the mechanism of heme-mediated platelet mtROS production which is dependent on the activation of platelet TLR4 signaling and leads to the downstream phosphorylation of complex-V by the serine kinase Akt. Notably, inhibition of platelet TLR4 or Akt, or scavenging mtROS prevents heme-induced granule release in vitro. Further, heme-dependent granule release is significantly attenuated in vivo in mice lacking TLR4 or those treated with the mtROS scavenger MitoTEMPO. These data elucidate a novel mechanism of TLR4-mediated mitochondrial regulation, establish the mechanistic link between hemolysis and platelet degranulation, and begin to define the heme and mtROS-dependent platelet secretome. These data have implications for hemolysis-induced thrombo-inflammatory signaling and for the consideration of platelet mitochondria as a therapeutic target in hemolytic disorders.

**Key points:**

1. Heme induces platelet mtROS production by inhibiting complex-V activity via TLR4 signaling.
2. Heme stimulated platelet granule secretion is regulated by mtROS.

## Introduction

Intravascular hemolysis occurs in a number of pathologies ranging from genetic hemoglobinopathies^1–3^ to more acute conditions such as sepsis^4^, pre-eclampsia^5,6^, parasitic infection^7,8^, or in patients after cardiac surgery^9,10^. It is well established that patients with chronic hemolysis, such as in sickle cell disease, are at significantly greater risk for thrombotic complications^11,12^ as well as endothelial dysfunction^13,14^ and chronic vasculopathy^15–17^ which lead to morbidities such as stroke^18,19^ and pulmonary hypertension (PH)^20–24^. However, the mechanisms that link hemolysis, thrombosis, and vasculopathy have not fully been elucidated. Platelets, when activated, are central mediators of thrombosis^25–27^, sentinels of inflammatory signaling ^27,28^, and can also propagate vascular responses through the synthesis and release of vasoactive molecules from alpha and dense granules^29–31^. Notably, cell-free hemoglobin (Hb) or free heme released via hemolysis stimulates platelet thrombotic activation^32–36^ and promotes inflammatory signaling^37,38^. At a mechanistic level, these effects have been linked to Hb-dependent modulation of platelet mitochondrial function. Specifically, Hb inhibits complex V of the platelet mitochondrial electron transport chain, leading to an increase in mitochondrial reactive oxygen species (mtROS) production, which stimulates platelet activation^32^. Consistent with this mechanism, specific scavengers of mtROS attenuate hemolysis-induced platelet activation^32^ and inflammatory signaling^39^ ex vivo and thrombosis in murine models^40^. Despite the recognition that Hb-induced mtROS regulates platelet activation and inflammatory signaling, the mechanisms by which hemoglobin or heme released via hemolysis stimulates mtROS production within the platelet is unknown.

Platelet degranulation, while associated with platelet activation, is a tightly regulated process. Although it is estimated that platelet granules contain over three hundred diverse molecules^41,42^, specific patterns of granule contents are released in response to differential platelet agonists^31,43–45^. Thrombospondin-1 (TSP-1) a multifunctional glycoprotein that is stored in and secreted from platelet α-granules regulates thrombosis^46–48^ as well as inflammatory^49,50^ and vascular signaling^50,51^. For example, once secreted, TSP-1 interacts with cell adhesive receptors and integrins to potentiate platelet activation and stabilize platelet aggregation^47,52,53^. Through its selective interaction with CD36 or CD47 on macrophages, other leukocytes, and endothelial cells, TSP-1 can stimulate the inflammatory response through the potentiation of NFkB^54^ and TGF-β signaling^49,55,56^ and enhancement of leukocyte migration^49,57^. Thrombospondin-1 also inhibits endothelial nitric oxide signaling and enhances matricellular remodeling to propagate vascular remodeling^58,59^. Notably, plasma and platelet TSP-1 levels are significantly increased in conditions with components of hemolysis such as sickle cell disease^60,61^ and sepsis^62,63^ respectively, and genetic inhibition of TSP-1 signaling attenuates vasculopathy in murine models of sickle cell disease^64^ and prevents inflammation and morbidity in a murine model of cecal ligation and puncture induced sepsis^65^. While these studies highlight the role of platelet-derived TSP-1 as a mediator of thrombo-inflammation and vasculopathy, it is unknown whether hemolysis-derived products stimulate TSP1 release, what other granule molecules are released by platelets upon encountering heme, and whether heme-dependent degranulation is regulated by platelet mtROS production.

In this study, we test whether heme and Hb released via hemolysis stimulate platelet granule release and determined the role of mtROS in this process. We demonstrate that cell free heme is a more potent stimulator of platelet mtROS production and TSP-1 release than Hb, and that mtROS production is required for the heme-dependent release of TSP-1 and other granule molecules. Further, we demonstrate that mechanistically heme-induced mtROS production requires the activation of platelet TLR4 signaling culminating in the activation of the serine/threonine kinase Akt, which phosphorylates complex V to inhibit its activity, leading to mtROS generation. These data have implications for the regulation of hemolysis induced thrombotic and vascular signaling, as well as for platelet mitochondria as a therapeutic target in hemolytic disease.

## Materials and Methods

All Chemicals were purchased from Sigma-Aldrich (St. Louis, MO) and antibodies from BD Biosciences (San Jose, CA) unless otherwise noted.

### Human blood collection and platelet isolation

Venous blood was collected from human participants by standard venipuncture after written informed consent was obtained and in accordance with study #19030018 (approved by the Institutional Review Board of the University of Pittsburgh).

Platelet rich plasma (PRP) was separated from whole blood collected in acid-citrate dextrose (ACD) Solution-A anticoagulant by centrifugation at 500 rpm for 20 minutes. Platelets were then pelleted in the presence of PGI_2_ (1μg/mL) by centrifuging the PRP at 1500xg for 10 minutes. These platelets were washed with erythrocyte lysis buffer containing PGI_2_ to remove any residual erythrocytes and resuspended in modified Tyrode’s buffer (20 mM HEPES, 128 mM NaCl, 12 mM sodium bicarbonate, 0.4 mM sodium phosphate monobasic, 5 mM dextrose, 1 mM MgCl_2_, 2.8 mM KCl, pH 7.4). The platelet count was determined by Hemavet® 950.

### Murine Studies

TLR4-flox and global TLR4 knockout mice (TLR4^−/−^) were used in accordance with approval from the University of Pittsburgh Institutional Animal Care and Use Committee. Male mice 10-12 weeks in age (24-27g) were administered cell free heme (110 mg/kg) or saline (vehicle) by tail vein injection. Some groups of mice were pre-treated with MitoTEMPO (300 μM) administered in the drinking water for 72 hours. Platelet mtROS (assessed via mitoSOX as described below) and cell free plasma concentrations of TSP-1, interleukin-1 beta (IL-1β), and platelet derived growth factor-B (PDGF-B) were measured 20 min after heme administration.

### Measurement of Platelet activation

Platelet activation was measured in washed platelets by staining with anti-CD41a-PE, anti-CD62P (P-selectin)-APC and PAC1 binding antibody-FITC (to bind activated GPIIb/IIIa), and then quantification of these markers and CD41a (as a marker of platelets) by flow cytometry (LSR-Fortessa; Becton Dickinson).

In all experiments, washed platelets (2.0-2.5 × 10^6^/mL) were incubated with heme for 30 minutes prior to measurement of activation. In some experiments, platelets were pre-treated with TLR4 neutralizing antibody (5 μg/mL), 2 μM of BX795 (TBK1 inhibitor), 100 nM of N-[1-[2-(4-Morpholinyl) ethyl]-1H-benzimidazol-2-yl]-3-nitrobenzamide (IRAK1/4 inhibitor), 5 nM of (5Z)-7-Oxozeaenol (TAK1 inhibitor), and/or 10μM of MitoTEMPO (mtROS scavenger).

### Measurement of mtROS

Treated washed platelets were pelleted at 1500xg for 5 min, resuspended in HBSS and incubated with 10 μM of MitoSOX™ Red for 5 min, after which fluorescent intensity (510/580 nm) was measured kinetically as previously described^32^. During the pretreatment of platelets with ARQ092 DMSO was used as vehicle control.

### Measurement of Thrombospondin-1 (TSP-1) release from platelets

TSP1 levels were measured using the Human Thrombospondin-1 DuoSet ELISA kit (R&D systems; DY3074) in the supernatant surrounding treated platelets. During the pretreatment of platelets with ARQ092 DMSO was used as vehicle control.

### Dot blot assay

Conditioned supernatant collected from treated washed platelets was blotted (30μL) on to the 0.4μm nitrocellulose membrane using Bio-Dot Apparatus. The membrane was blocked with blocking buffer for 30 min at room temperature and followed by incubation with primary antibody overnight at 4°C (all primary antibodies used for dot blot were purchased form, R&D systems; anti-Cathepsin A, AF1049; anti-Angiostatin, AF226; anti-CD40L, AF617; anti-Kininogen, AF1569; anti-PAI-1, AF1786; anti-Thrombospondin-1, AF3074; AF795; anti-CXCL7, AF393; anti-IL-1β, AF201; anti-PDGF-B, AF220; anti-TGF- β, AF246; anti-FGF basic, AF233) and IRDye® 800CW Donkey anti-Goat IgG secondary antibody for 40min at room temperature. The blots were imaged using a LI-COR imaging system and analyzed using Image Studio software.

### Mitochondrial Complex V activity assay

The enzymatic activity of complex V was measured by spectrophotometrically by kinetic assay as previously described^32^.

### Immunoprecipitation of Complex V

Treated platelets were lysed with RIPA lysis buffer including Halt protease and phosphatase inhibitor cocktail. The beta subunit of complex V and proteins bound to it were immunoprecipitated using an antibody to the beta subunit of ATP synthase (Millipore, MAB3494) in accordance with the Pierce™ Co-Immunoprecipitation Kit (Cat.No:26149). Immunoprecipitated samples were subjected to western blot by standard procedure using anti-pAKT (S473; Cell Signaling, 4060S) and anti-ATP synthase beta subunit antibodies. The blots were imaged using a LI-COR imaging system and analyzed using Image Studio software.

### Statistics

Unpaired parametric t-tests were used to compare individual group samples and ANOVA along with Tukey post-hoc test were used to make multiple comparisons. Statistical analyses were performed using GraphPad Prism 9 software. P-values <0.05 were considered significant. Data are presented as mean ± standard error of the mean (SEM) unless otherwise specified.

## Results

### Heme is a more potent platelet agonist than Hb

To determine whether heme stimulates platelet activation as potently as Hb, isolated washed human platelets were incubated with heme (0-20μM) or Hb (0-40μM) and platelet activation was measured. Platelets treated with heme or Hb both showed a concentration dependent increase in surface P-selectin levels and PAC-1 antibody binding (as a measure of activated GPIIb/IIIa), indicative of platelet activation **(Figure 1A-B)**. However, heme treatment stimulated a greater level of surface P-selectin and active GPIIb/IIIa at every concentration measured (70.9 ± 4.03% P-selectin; 68.14 ± 2.76% PAC-1 binding at 2.5 μM) compared to Hb (7.59 ± 0.40% P-selectin; 10.41 ± 2.24% PAC-1 binding at 2.5 μM) **(Figure 1A-B)**. These results indicate that heme and Hb both activate platelets, but heme is a more potent agonist than Hb for platelet activation. To investigate the role of heme in modulating platelet function beyond activation, we measured the release of TSP-1 from α-granules of heme or Hb treated platelets, as a marker of platelet granule secretion. We observed significantly greater levels of TSP-1 release in heme stimulated platelets compared to those stimulated with Hb **(Figure 1C),** similar to the effect on platelet activation. Since the platelet granule secretome is agonist specific, we next measured the level of release of a panel of nine common platelet granule molecules in response to heme stimulation. We found that heme induced the release of seven of the granule factors measured (CXCL7, FGF basic, TGFβ, IL-1β, PDGF-B, angiostatin, kininogen), while it did not stimulate the release of CD40L and PAI-1 (**Figure 1D**). These data demonstrate that heme stimulates the release of a specific pattern of granule molecules.

**Figure 1.**
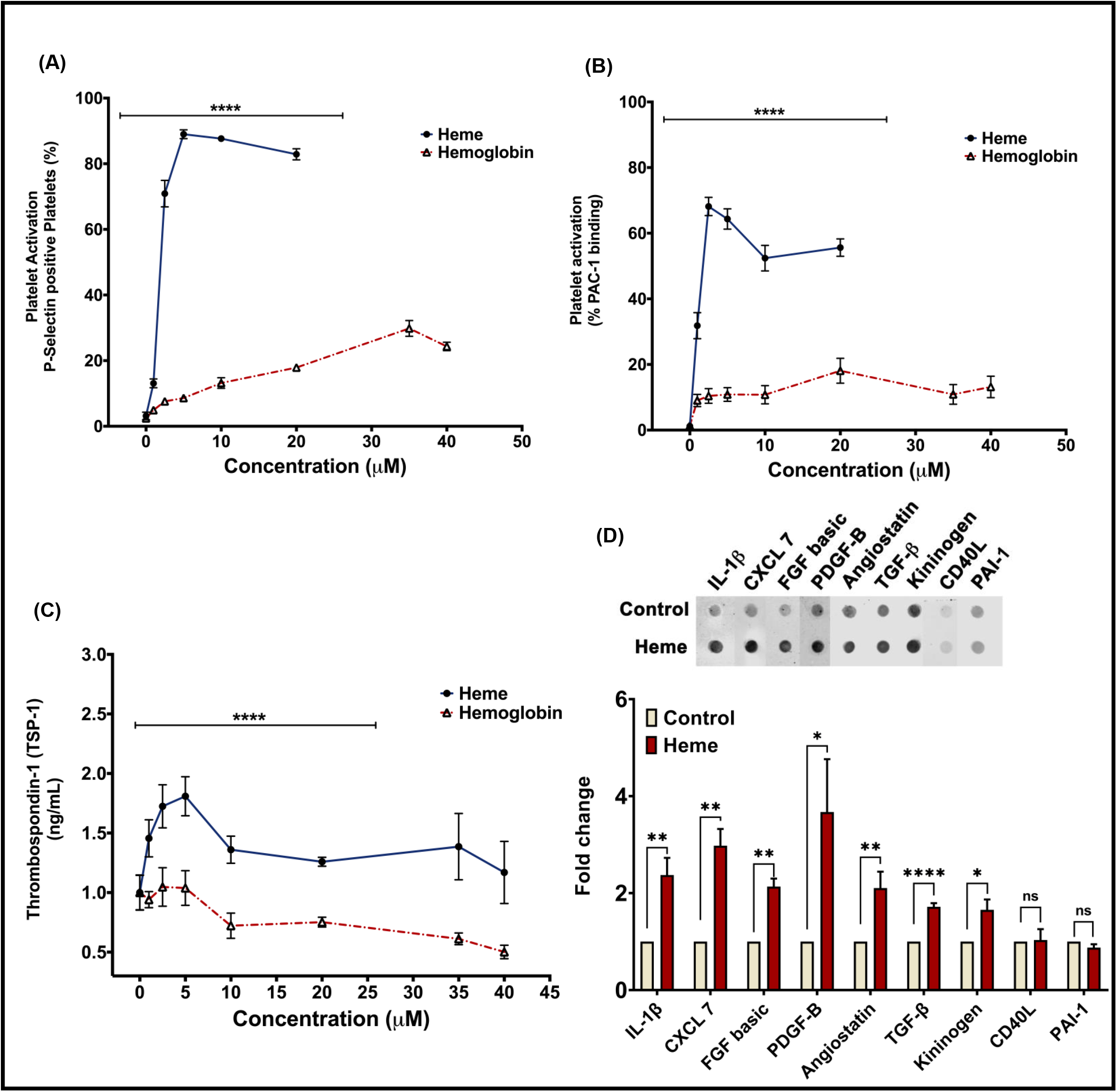
Heme is a more potent platelet agonist than hemoglobin. Platelet activation measured by **(A)** platelet surface P-selectin or **(B)** activated GPIIb/IIIa levels in heme (blue line) or methemoglobin (red line) treated platelets. **(C)** Thrombospondin-1 levels quantified in the releasate from heme (blue line) or methemoglobin (red line) treated platelets. **(D)** Dot blot with quantification of IL-1β, CXCL7, FGF basic, PDGF-B, angiostatin, TGFβ, kininogen, CD40L, PAI-1 levels in the releasate from heme (2.5μM) treated platelets. Data are represented as Mean ± SEM. \*\*\*\**p*<0.0001, \*\**p*<0.01, \**p*<0.05, ns- not significant. n=3.

### Heme- induced mtROS production stimulates platelet granule release

We previously showed that Hb inhibits platelet mitochondrial complex V activity, which increases mitochondrial inner membrane potential to promote mtROS production, an effect that was associated with platelet activation^32^. To test whether heme similarly modulates platelet mitochondrial function, we treated platelets with heme and measured complex V activity and mtROS production. Heme treatment significantly decreased platelet mitochondrial complex V activity **(Figure 2A),** and concomitantly increased mtROS production **(Figure 2B)**.

**Figure 2.**
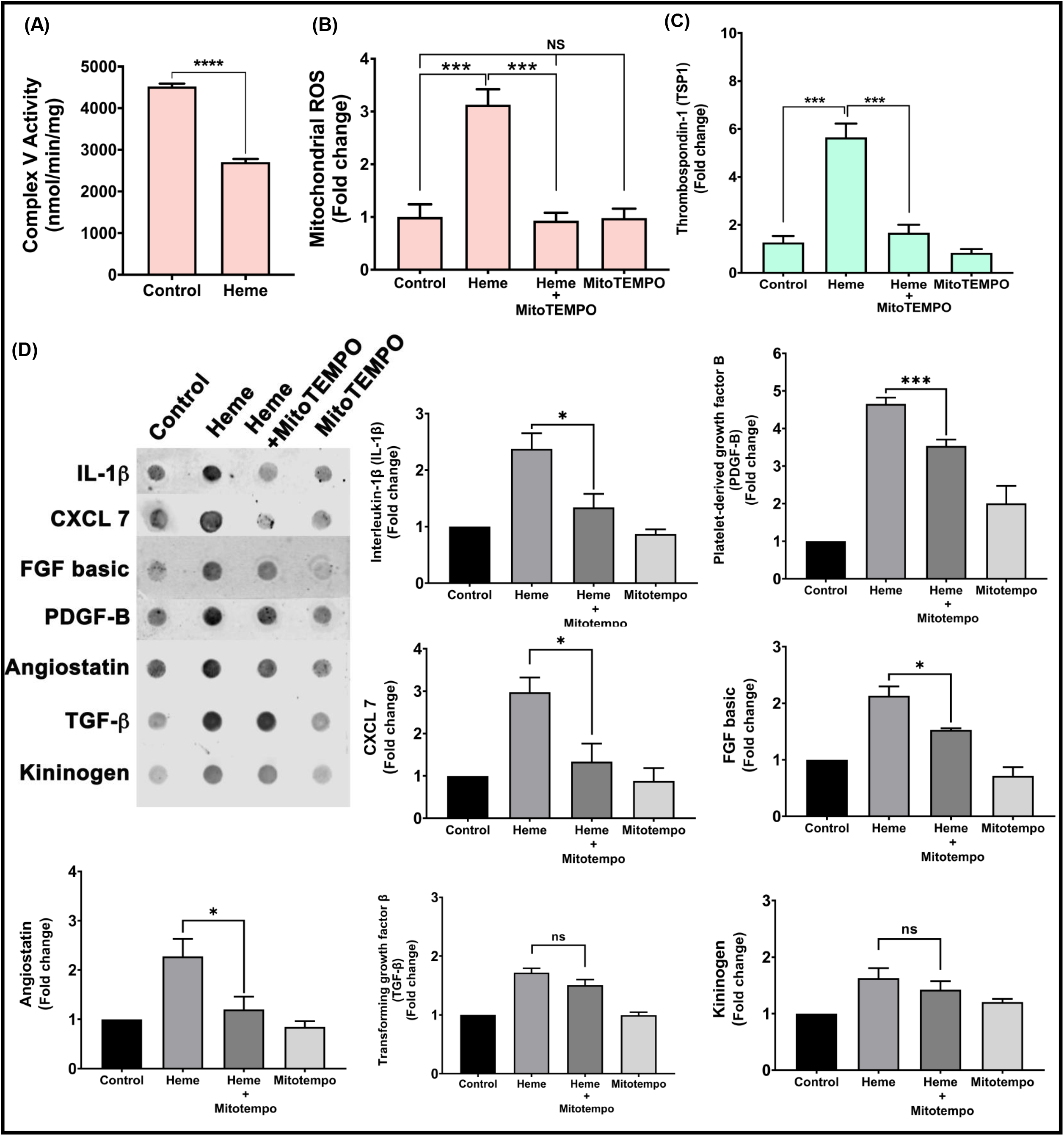
Heme inhibits platelet complex V activity and induces mtROS production that stimulates granule release. Platelets were treated with heme (2.5 μM) in the presence or absence of MitoTEMPO (10 μM) or with mitoTEMPO alone and **(A)** Platelet mitochondrial complex V activity and **(B)** mtROS production were measured. **(C)** Thrombospondin-1 levels in the platelet releasate were measured. **(D)** Dot blot along with the quantification of levels of IL-1β, CXCL7, FGF basic, PDGF-B, angiostatin, TGFβ and kininogen in the platelet releasate. Data are Mean ± SEM. \*\*\*\**p*<0.0001, \*\*\**p*<0.001, \**p*<0.05, ns- not significant. n=4

To determine whether mtROS production was required for platelet granule release, we measured the levels of release of the eight granule factors identified to be stimulated by heme (TSP1, CXCL7, FGF basic, TGFβ, IL-1β, PDGF-B, angiostatin, kininogen) from heme treated platelets in the presence and absence of MitoTEMPO (10μM), a mtROS scavenger. Treatment of platelets with MitoTEMPO significantly decreased heme-induced mtROS levels **(Figure 2B)** and also significantly attenuated heme-induced release of TSP1, CXCL7, FGF basic, IL-1β, PDGF-B, angiostatin **(Figure 2C-D)**.

### Heme inhibits mitochondrial complex V and induces mtROS in a TLR4 dependent manner

To determine the mechanism by which extracellular heme mediates intra-platelet signaling, we tested whether a platelet surface receptor was required for heme-mediated platelet granule secretion, focusing on TSP-1 as a marker of heme induced granule release. Since TLR4 is known to mediate heme-dependent responses in other cell types^66,67^, we blocked platelet TLR4 with TLR4 neutralizing antibody (5 μg/mL) and measured complex V activity and mtROS production in platelets after treatment with heme (2.5 μM). Blocking platelet TLR4 attenuated heme-dependent complex V inhibition **(Figure 3A)** and significantly decreased heme-induced mtROS production **(Figure 3B)**. Consistent with heme induced TSP-1 secretion being dependent on mtROS production, the presence of TLR4 neutralizing antibody also significantly decreased heme-induced TSP-1 secretion **(Figure 3C).**

**Figure 3.**
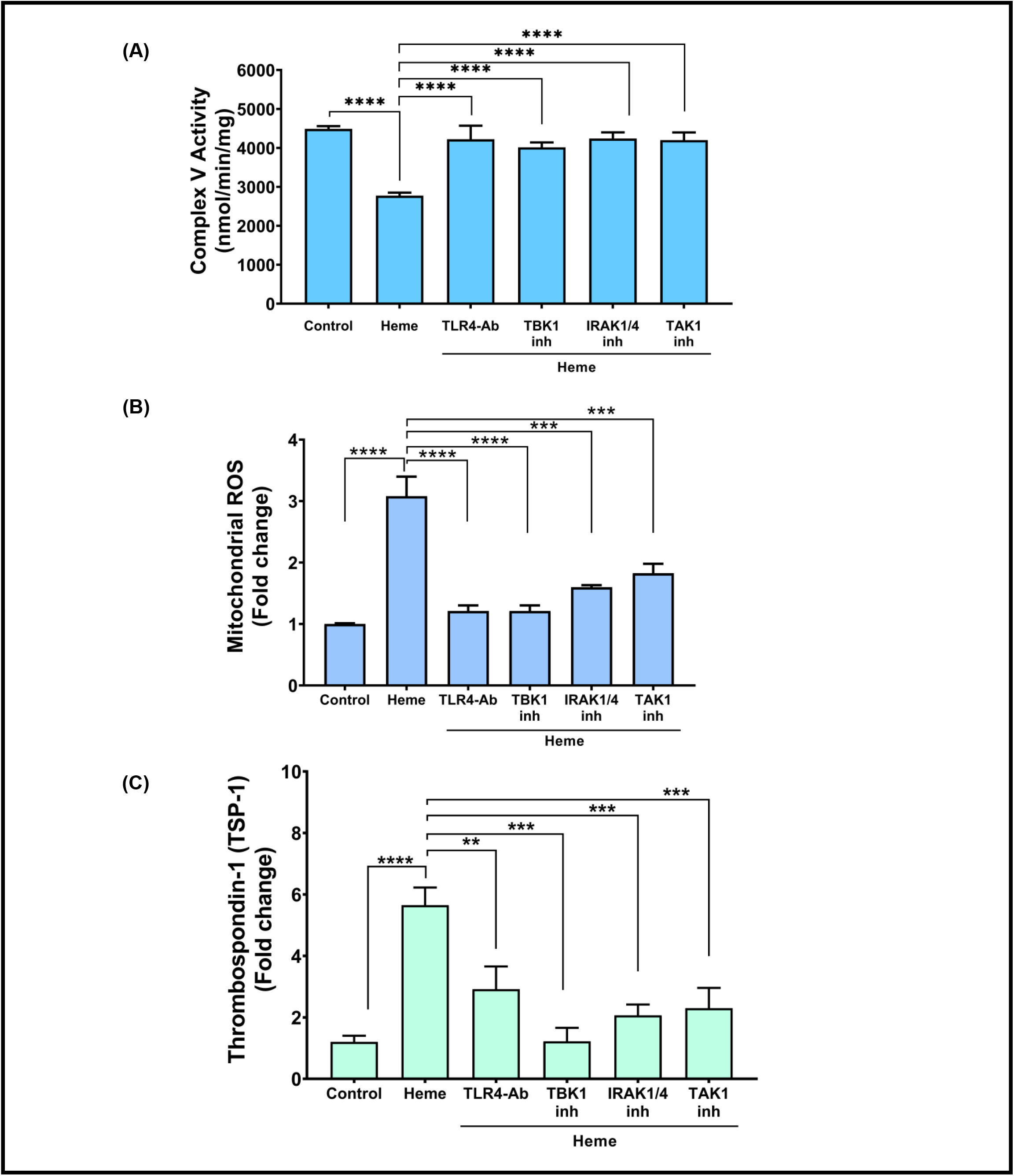
Heme inhibits platelet complex V activity to induces mtROS production and TSP-1 release via TLR4. **(A)** Platelet mitochondrial complex V activity, **(B)** platelet mtROS production, and **(C)** thrombospondin-1 levels in the releasate were measured in heme-treated platelets in the presence or absence of TLR4 neutralizing antibody (5 μg/mL), pharmacological inhibitors of TBK1 (2 μM of BX795), IRAK1/4 (100 nM of N-[1-[2-(4-Morpholinyl)ethyl]-1H-benzimidazol-2-yl]-3-nitrobenzamide), TAK1 ( 5 nM of (5Z)-7-Oxozeaenol). Data are represented as Mean ± SEM. \*\*\*\**p*<0.0001, \*\**p*<0.01, \*\*\**p*<0.001. n=4

Given that blocking platelet surface TLR4 resulted in decreased heme-induced platelet mtROS production by improving complex V activity and attenuated TSP-1 secretion from heme treated platelets, we sought to determine how TLR4 downstream signaling inhibits complex V activity. We used pharmacological inhibitors to inhibit key downstream kinases in the TLR4 pathway. We blocked signaling associated with the TLR4 adaptor protein MyD88 with inhibitors of kinases downstream of MyD88 - IRAK1/4 and TAK1. Additionally, we used inhibitors of TBK1 kinase to test the role of MyD88-independent signaling. Inhibition of TBK1, IRAK1/4 or TAK1 individually significantly attenuated heme-dependent complex V inhibition **(Figure 3A)**. Similarly, mtROS production in heme-treated platelets was significantly decreased in the presence of the inhibitors of TBK1, IRAK1/4 or TAK1. **(Figure 3B)**. Consistent with the requirement for complex V inhibition and mtROS production to induce TSP-1 release by heme, inhibitors of downstream TLR4 signaling also significantly attenuated TSP-1 release from the heme-treated platelets **(Figure 3C).** Collectively, these data suggest that heme-dependent TLR4 activation inhibits mitochondrial complex V and induces mtROS production through MyD88 dependent and independent pathways.

### Heme mediated TLR4 signaling promotes AKT phosphorylation to inhibit complex V activity

Akt is a serine/threonine-specific protein kinase that can be activated downstream of TLR4 and is known to bind to and phosphorylate a number of mitochondrial proteins, including the α and β subunits of complex V, to regulate their function ^68,69^. To determine whether heme-induced TLR4 activation requires AKT activation to inhibit complex V activity, we tested whether AKT binds to complex V and examined the phosphorylation status of AKT in heme treated platelets. Immunoprecipitation of the beta-subunit of complex V from heme treated platelets showed that significant levels of Akt were associated with complex V, and measurement of pAKT(S473) showed significant activation of the kinase **(Figure 4A)**. When phosphorylation of AKT was assessed in heme treated platelets pre-treated with blockers of TLR4 signaling (TBK1, IRAK1/4 or TAK1 inhibitors), lower levels of pAKT(S473) bound to complex V were measured **(Figure 4A)**. These data demonstrate that heme-mediated activation of TLR4 stimulates downstream AKT phosphorylation and binding to complex V.

**Figure 4.**
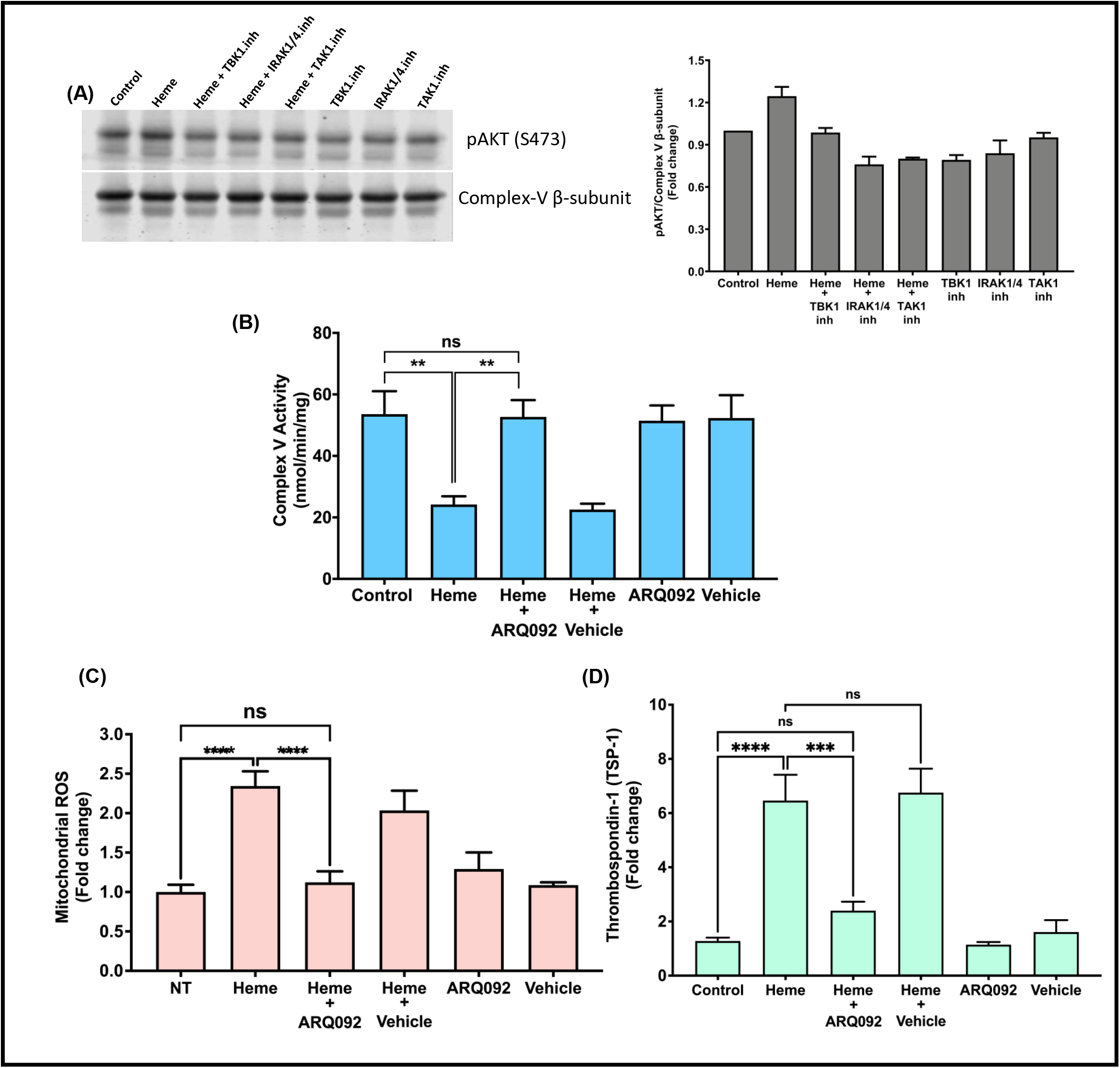
Heme-induced TLR4 signaling activates AKT to inhibit complex V activity in platelets. **(A)** Platelets were treated with heme (2.5 μM) in the presence or absence of TLR4 downstream signaling blockers (2 μM of BX795 (TBK1.inh), 100 nM of N-[1-[2-(4-Morpholinyl)ethyl]-1H-benzimidazol-2-yl]-3-nitrobenzamide (IRAK1/4.inh) or 5 nM of (5Z)-7-Oxozeaenol (TAK1.inh), and pAKT(S473) was measured in immunoprecipitated Complex-V beta subunit from platelet lysates. pAKT(S473) levels were normalized with complex V beta subunit and plotted as fold change. **(B)** Platelet mitochondrial complex V activity **(C)** mtROS production and **(D)** thrombospondin-1 release were measured in platelets pre-treated with or without ARQ092 (10 μM) treated with heme (2.5 μM). Data are represented as Mean ± SEM. \*\*\*\**p*<0.0001, \*\*\**p*<0.001, \*\**p*<0.01, ns – not significant. n=4.

To determine whether heme-induced activation of AKT regulates complex V activity, we treated platelets with heme in the presence and absence of ARC0092, a small molecule that prevents phosphorylation of AKT at S473^70^, and measured complex V activity. While heme treatment inhibited complex V activity, this effect was significantly attenuated when AKT phosphorylation was blocked **(Figure 4B)**. Collectively, these data demonstrate that heme-induced TLR4 activation stimulates the downstream activation of AKT, which binds to complex V and inhibits its activity.

Given that heme induced mtROS production is essential for TSP-1 release from platelets, and complex V inhibition (which leads to mtROS production) is dependent on AKT phosphorylation, we tested whether blocking AKT activation decreases heme-induced mtROS production and TSP-1 release. We pretreated platelets with ARC0092, incubated them with heme and measured platelet mtROS production and TSP-1 release. As expected, heme increased both platelet mtROS production and TSP-1 release. However, both these effects were significantly decreased upon blocking AKT phosphorylation **(Figure 4C-D)**.

To determine whether the heme-induced pathway elucidated ex vivo was relevant in a physiological setting in vivo, TLR4^−/−^ mice and corresponding control mice (TLR4 flox) were administered cell free heme (110mg/kg) to mimic hemolysis and platelet mtROS was measured. While heme induced platelet mtROS production in control mice, this effect was significantly attenuated in TLR4^−/−^ mice (**Figure 5A**). Measurement of plasma levels of TSP-1, PDGF-B, and IL-1β showed that heme-induced release of these factors was also significantly attenuated in TLR4^−/−^ mice. Consistent with the dependence of mtROS on TLR4 signaling, treatment with MitoTEMPO attenuated heme-induced granule secretion in control mice but had no effect in TLR4^−/−^ mice (**Figure 5B**).

**Figure 5.**
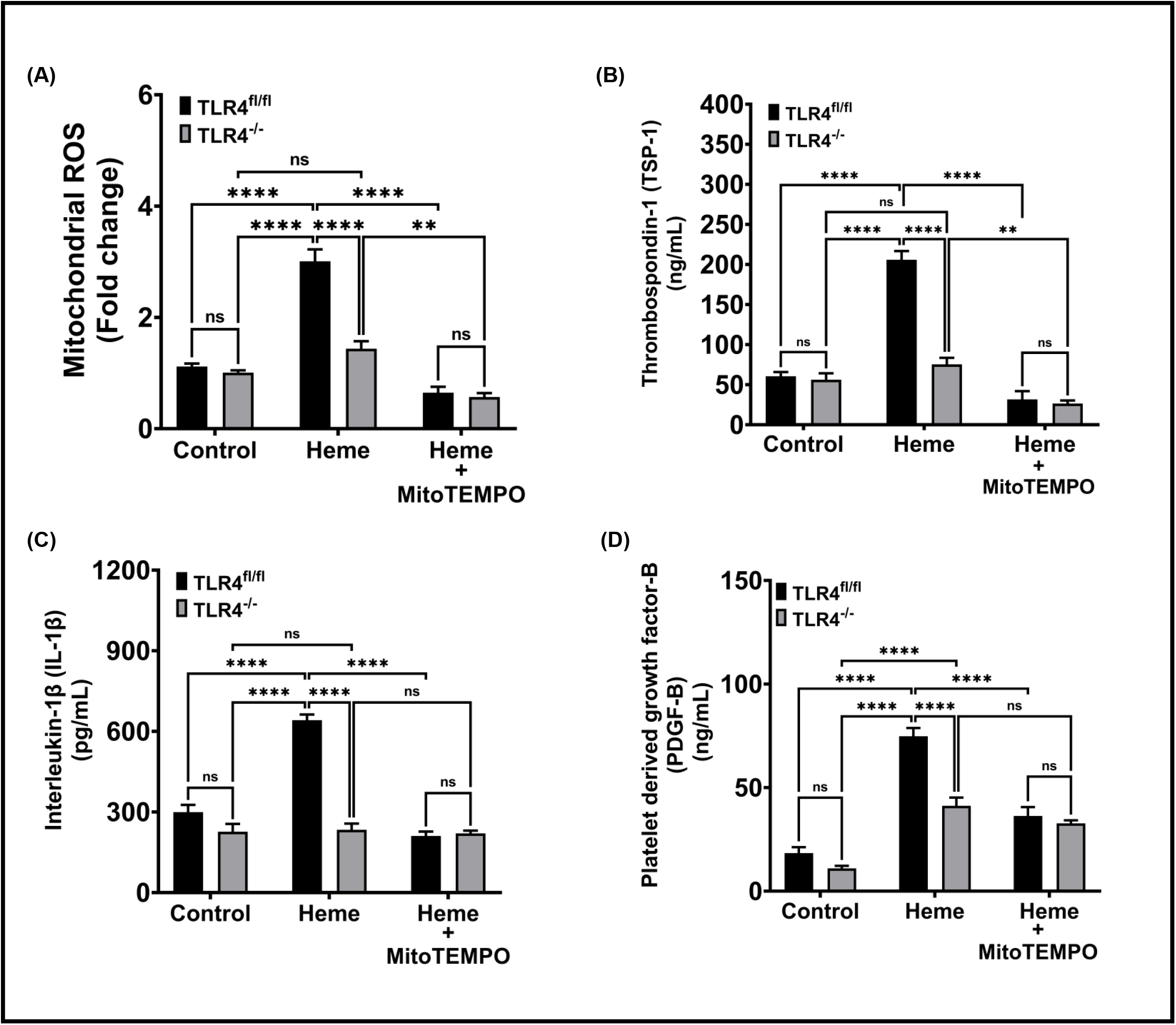
Heme infused TLR4^−/−^ mice show attenuated mtROS production and heme induced granule release in platelets. TLR4^−/−^ mice and TLR4^fl/fl^ mice (control mice) were pretreated with MitoTEMPO (300μM) and injected (i.v.) with heme (110mg/kg). (A) Platelets mtROS was measured and plasma levels of (B) Thrombospondin-1, (C) Interleukin-1β and (D) Platelet derived growth factor-B were measured. Data are represented as Mean ± SEM. *\*\*\*\*p<0.0001, **p<0.001, ns- not significant. n=5 mice per group in control and heme treated* groups; n=3 to 4 mice in MitoTEMPO treated groups.

## Discussion

In this study we demonstrate that while heme and hemoglobin both stimulate platelet activation and TSP-1 release, heme is a more potent platelet agonist than Hb. Mechanistically, we show that heme-mediated TSP-1 release relies on the production of platelet mtROS generation. Further, we elucidate the mechanism by which heme stimulates mtROS production, and find that mtROS production is dependent on heme-mediated activation of TLR4 signaling, culminating in the Akt-dependent phosphorylation and inhibition of complex V activity. This pathway has implications for not only understanding the pathogenesis of hemolysis-induced thrombotic and inflammatory diseases, but also for the development of potential therapeutics for these conditions.

Prior studies have independently reported that heme^35,36^ and Hb^32–34^ stimulate platelet activation, however their potency in the context of platelet agonism has previously not been compared. We show here that heme is a more potent mediator of both platelet activation and TSP-1 release than Hb. It is unclear what factors underlie this increased potency of heme. Heme and Hb are biochemically distinct species and heme can selectively bind to multiple receptors and transcription factors^35,67,71,72^. Thus, it is likely that heme is either a stronger activator of TLR4 or stimulates multiple signaling pathways that concomitantly contribute to platelet activation and/or TSP-1 secretion. While prior studies demonstrate heme-induced activation of TLR4^67,71^, it remains unclear whether heme activates TLR4 through traditional ligand binding, and whether Hb acts similarly. However, consistent with the potential of heme activating concomitant pathways of platelet agonism, Bourne and colleagues showed that heme-mediated activation of C-type-lectin-like-receptor-2 (CLEC-2) contributes to platelet activation^35^. Notably, CLEC-2 activation was observed with higher concentrations of heme (6.25 μM)^35^ than used in our study (2.5 μM). Given the biphasic effect of heme observed by both groups, in which maximal platelet agonism is observed at ~5-6μM heme, it is possible that while both pathways contribute to heme-mediated platelet agonism, the contribution of these pathways potentially shifts with increasing concentrations of heme. Further study is required to delineate the contribution of each pathway, their potential cross-talk, and whether stimulation of multiple pathways makes heme a more efficient platelet agonist than Hb.

Though heme-mediated platelet activation is well documented ^35,36^, heme-induced platelet granule release has not been extensively studied. It is estimated that platelet granules store over three hundred molecules, including mitogens, chemokines, and thrombotic regulators, which are released in discreet patterns dependent on the specific stimulus^31,43–45^. This study begins to define the heme-dependent platelet secretome by showing that heme induces the release of TSP-1, PDGF-B, IL-1β, FGF basic, angiostatin and CXCL7. Importantly, we also identified molecules that are not released in response to heme such as CD40L and PAI-1. Identification of the heme specific platelet secretome may provide a critical link between hemolysis and pathogenic vascular signaling. For example, platelet derived TSP-1 can promote both thrombosis and vascular pathogenesis through its interaction with CD36 on circulating cells and CD47 in endothelial cells ^47,73–75^. In severe sepsis patients, plasma TSP-1 levels have been found to be significantly elevated^63^, and recent murine studies of sepsis models show that independent of pathogen load, free heme promotes thrombosis^76^. Thus, it is interesting to speculate that heme-induced platelet TSP-1 release potentially drives pathogenic thrombosis in sepsis. Similarly, in sickle cell disease, platelet TSP-1 has been implicated in the pathogenesis of pulmonary hypertension^77^, which is a major cause of morbidity in these patients and is also associated with hemolysis^60^. Notably, plasma TSP-1 levels are significantly elevated in patients with sickle cell disease in steady state^60,78^. In patients who are in vaso-occlusive crisis, which is associated with even higher rates of hemolysis than in steady state, TSP-1 levels associate with lower rates of hemolysis^60^. This is likely consistent with the biphasic curve (Figure 1) for heme-dependent TSP-1 release that we demonstrate in this study.

While prior studies have demonstrated that heme induces ROS production in the platelet, this study is the first to demonstrate the mitochondrion as a significant source of heme-induced ROS and to define the mechanism by which heme induces mtROS. Our data demonstrate that heme inhibits mitochondrial complex V to induce mtROS production, and that this inhibition of complex V requires heme-mediated activation of platelet TLR4 signaling which ultimately results in Akt activation and association with complex V. Our data are consistent with accumulating reports demonstrating that phosphorylated Akt can translocate and accumulate in the mitochondria, and specifically that Akt can phosphorylate mitochondrial complexes, including the β-subunit of complex-V^68,69^. Our data are not entirely consistent with other reports of Akt-dependent complex V phosphorylation which show that this association leads to activation of the complex V rather than the inhibition shown in this study. However, inhibitory and activating phosphorylation sites have been identified on complex V β-subunit. Thus, it is possible that separate stimuli propagate Akt-dependent phosphorylation of different sites. Further study identifying the heme-dependent phosphorylation site is required for comparison with other stimuli.

The data presented in this study demonstrate that mtROS regulate heme-dependent granule release. While a growing number of reports have established the association between mtROS production and platelet activation in multiple pathologies^79–81^, the specific role of mtROS in regulating platelet function, particularly granule secretion in response to specific agonists, is less clear. Notably, a recent study demonstrates that agonists such as thrombin receptor activator peptide-6 (TRAP-6) activates platelets in a mtROS independent manner, and while scavenging mtROS does not affect TRAP-6 dependent platelet activation, it significantly attenuates platelet aggregation^80^. These data are consistent with mtROS regulation of platelet granule secretion independent of regulation of platelet activation. However, further study is required to dissect mtROS regulation of platelet activation versus granule secretion, as well as the molecular mechanisms by which mtROS cause granule release. However, our in vivo data suggest that mtROS scavenging (or TLR4 inhibition to prevent mtROS production) is a potentially viable option in preventing heme-induced platelet dysfunction.

In conclusion, this study demonstrates the mechanism by which extracellular heme signals through platelet TLR4 to induce platelet mtROS production. Further, we demonstrate that heme-induced mtROS stimulates platelet granule secretion. The data begin to define the heme-induced platelet secretome and its regulation by mtROS. Overall, these studies advance the understanding of the mechanisms that link hemolysis to platelet dysfunction. While further study of the secretome is required to determine whether heme-induced secreted products are responsible for hemolysis-associated inflammation and vasculopathy, the data herein demonstrate a central role for the mitochondrion in heme-dependent platelet dysfunction and suggest this organelle as a potential therapeutic target in hemolytic conditions.

## Acknowledgments

This work was supported by NIH grants HL133003-01A1 (to SS) and 1 R01 HL130268-03 (to SS and BZ) and The Hemophilia Center of Western Pennsylvania (to SS).

## Authorship Contributions

GA, DND, MR, YW, LK, BZ and SS designed the study and performed the research. LK and BZ performed murine experiments and collected the data.GA, DND, MR and SS analyzed the data. GA and SS wrote the manuscript.

## Conflict of Interest Disclosures

The authors declare no competing financial interests.

## Notes

### Competing Interest Statement

The authors have declared no competing interest.

